# Norway spruce deploys tissue specific canonical responses to acclimate to cold

**DOI:** 10.1101/2020.01.13.904805

**Authors:** Alexander Vergara, Julia Christa Haas, Paulina Stachula, Nathaniel Robert Street, Vaughan Hurry

## Abstract

Cold acclimation in plants is a complex phenomenon involving numerous stress-responsive transcriptional and metabolic pathways. Existing gene expression studies have primarily addressed short-term cold acclimation responses in herbaceous plants, while few have focused on perennial evergreens, such as conifers, that survive extremely low temperatures during winter. To characterize the transcriptome changes during cold acclimation in *Picea abies* (L.) H. Karst (Norway spruce), we performed RNA-Sequencing analysis of needles and roots subjected to a chilling progression (5 °C) followed by 10 days at freezing temperature (−5 °C). Comparing gene expression responses of needles against *Arabidopsis thaliana* L. (Arabidopsis) leaves, our results showed that early transient inductions were observed in both species but the transcriptional response of Norway spruce was delayed. Our results indicate that, similar to herbaceous species, Norway spruce principally utilizes early response transcription factors (TFs) that belong to the <u>AP</u>ETALA <u>2</u>/<u>e</u>thylene-<u>r</u>esponsive element binding factor (AP2/ERF) superfamily and NACs. However, unique to the Norway spruce response was a large group of TFs that mounted a late transcriptional response to low temperature. A predicted regulatory network analysis identified key conserved TFs, including a root-specific *bHLH101* homolog and other members of the same family with a pervasive role in cold regulation, such as homologs of *ICE1* and *AKS3* and also homologs of the NAC (*anac47* and *anac28*) and AP2/ERF superfamilies (*DREB2* and *ERF3*), providing new functional insights into cold stress response strategies in Norway spruce.

**One sentence summary:** Norway spruce shares elements of the cold regulon described in herbaceous species but has undescribed components that contribute to the cold tolerance of this evergreen coniferous species.

## Introduction

Plants vary in their capacity to tolerate cold stress, and low temperatures strongly limit species distribution and plant productivity (Levitt, 1980). Temperate herbaceous crop and model plant species, such as *Arabidopsis thaliana* L. (Arabidopsis), acclimate to cold temperatures after exposure to chilling, non-freezing temperatures (0-15 °C) and increase the freezing tolerance after exposure to temperatures below 0 °C (Miura and Furumoto, 2013). Studies on these species have provided insight into the complex molecular mechanisms involved and changes in global gene expression have been shown to support a multitude of metabolic and physiological modifications such as the accumulation of cryoprotective molecules (amino acids, amines, proteins and carbohydrates) and antioxidants, as well as adaptations in membrane fluidity in response to cold (Hurry et al., 1995; Murata and Los, 1997; Strand et al., 1999; Strand et al., 2003; Cook et al., 2004; Renaut et al., 2006; Janska et al., 2010; Hoermiller et al., 2016). The best studied of the cold response pathways is that regulated by the CBF/DREB1 (CRT-binding factor/DRE-binding protein) family of transcription factors (Shinozaki and Yamaguchi-Shinozaki, 1996; Fowler and Thomashow, 2002; Cook et al., 2004), which bind to a CRT/DRE (C-repeat/Dehydration Responsive Element) element in the promoter region of target genes (Stockinger et al., 1997). Investigations directed at improving frost tolerance in agricultural plants often target the CBF pathway because of its wide conservation in plants (Thomashow, 2010). However, in recent genome-wide transcriptomic analyses, other early cold-induced transcription factors and their role in regulating cold responsive (COR) genes have received attention, highlighting the need for whole-genome transcriptional analyses to unravel the complexity of the low-temperature gene regulatory network (Park et al., 2015).

In woody perennials, overwintering is initiated after sensing the change of season (Welling et al., 2004), with the shortening of the photoperiod in late summer/early autumn inducing growth cessation, the development of dormancy and cold hardening (Guy, 1990; Bigras et al., 2001; Li et al., 2004; Welling et al., 2004; Rossi et al., 2008; Cooke et al., 2012; Chang et al., 2015). However, maximal frost tolerance is only acquired after exposure to temperatures below 0 °C (Sakai, 1966; Weiser, 1970; Greer and Warrington, 1982; Bigras et al., 2001; Beck et al., 2004; Sogaard et al., 2009). The establishment of frost tolerance to extreme low temperatures (< −60 °C) in perennial or over-wintering tissues makes it possible for aboveground parts to survive above the snow cover and may be a result of greater cellular dehydration than found in herbaceous species (Lang et al., 1994; Welling et al., 1997; Rinne et al., 1998; Welling et al., 2004), coupled to more intensive accumulation of cryoprotective compounds (Coleman et al., 1991; Kuroda and Sagisaka, 1993; Rinne et al., 1994; Sauter and Wellenkamp, 1998). High concentrations of solutes also increase the intracellular viscosity, stabilizing cells when stressed and leading to the formation of aqueous glasses in woody plants (Wisniewski et al., 2003). Nevertheless, woody perennials and herbaceous plant species also share mechanisms of cold regulation and cold-regulated target genes. For example, CBFs have been found in deciduous (poplar, birch) (Nanjo et al., 2004; Benedict et al., 2006; Welling and Palva, 2008) and evergreen angiosperm tree species (eucalyptus) (El Kayal et al., 2006; Navarro et al., 2009) and are involved in the activation of the cold responses of leaves and winter dormant tissue, where specialization after perennial-driven evolution might explain differences in the transcriptomes.

Boreal forests cover about 11% of the earth’s surface (Bonan and Shugart, 1989) and are dominated by evergreen conifers of the genera *Abies*, *Picea* and *Pinus,* which can survive extended periods of temperatures below – 40 °C when fully cold acclimated (Sakai, 1966; Sakai and Weiser, 1973; Strimbeck et al., 2007, 2008). In contrast to deciduous trees of temperate and boreal regions, evergreen conifers such as Norway spruce (*Picea abies* (L.) H. Karst) maintain their photosynthetic tissues for several years. In order to keep the evergreen foliage alive throughout the winter the needles have to acquire extreme low-temperature tolerance (Strimbeck et al., 2007, 2008). Similar to angiosperms, changes in carbohydrates (Strimbeck et al., 2008), accumulation of low-molecular weight cryoprotectant metabolites (Crosatti et al., 2013; Chang et al., 2015) and cryoprotective proteins such as dehydrins (Kjellsen et al., 2013), contribute to acquired freezing tolerance in conifer needles.

Frost injuries can also occur in belowground tissues of plants, with soil frost causing fine root dieback, reducing nutrient and water uptake by trees (Groffman et al., 2001). Normally, snow cover insulates the soil from the cold air temperatures and reduces freeze-thaw events in the soil (Campbell et al., 2005), protecting roots from extreme temperature changes, and the question of whether and how roots develop cold tolerance has received little attention. However, Arabidopsis has been reported to show as little as 14% overlap in cold-induced leaf and root transcriptomes, based on 8K microarray chips (Kreps et al., 2002). Furthermore, the limited data available suggests that conifer roots remain metabolically active longer than above ground tissues and they do not develop the same deep frost tolerance as the above ground tissues (Bigras et al., 2001), suggesting that root responses to changing seasonal temperatures warrant closer investigation.

Climate change models predict that temperatures will increase and become more variable at higher latitudes, particularly during the winter months (Christensen et al., 2007). These changes in seasonal temperatures will not only increase the length of the growing season (Barichivich et al., 2013) but possibly also delay the onset of cold acclimation, impair the development of freezing tolerance in the autumn and lead to early deacclimation during the late winter (Repo et al., 1996; Wang, 1996; Guak et al., 1998; Stinziano et al., 2015; Chang et al., 2016; Frechette et al., 2016). Furthermore, current evidence also suggests that the risk of belowground frost injury is increasing (Campbell et al., 2005), with warmer winter air temperatures reducing the depth and duration of the snow cover, leading to colder soil temperatures (Groffman et al., 2001; Decker et al., 2003).

Conifers have evolved separately from angiosperms for more than 300 million years (Bowe et al., 2000) and large-scale whole-genome transcriptional profiling experiments are needed to understand whether the same gene regulatory networks are involved in cold acclimation in conifers. The assembly of a draft genome of Norway spruce (Nystedt et al., 2013) has made this species an ideal coniferous model. To gain insight into the mechanisms that have enabled conifers to dominate the boreal forest under current climatic conditions, we performed genome-wide RNA-Seq analysis from needles and roots of non-dormant two-year old Norway spruce seedlings exposed to cold (5 °C) and freezing (−5 °C) temperatures. This experimental design allowed us to focus only on the temperature responses, without influence from growth cessation and dormancy related mechanisms. For direct comparison with the standard herbaceous model, Arabidopsis leaves exposed to cold (5 °C) were sampled at equivalent time points.

## Results

### Comparing the cold acclimation transcriptional responses of Norway spruce needles and Arabidopsis leaves

RNA-Seq analysis of needles and leaves exposed to cold (5 °C) for different times revealed that the transcriptional response of Norway spruce needles was slower than for Arabidopsis leaves (Fig. 1A, B). Overall, both species differentially expressed approximately 15% of all transcribed genes in response to cold, with approximately equal numbers up- and down-regulated (Fig. 1B). When the homology relationships of the differentially expressed genes (DEGs) between species were compared using orthology information from the Gymno PLAZA resource (Van Bel et al., 2018) (Fig. 1C) it was found that of the 1949 Arabidopsis cold-induced DEGs, 340 had an identified ortholog that also responded to cold in Norway spruce (validated orthologs). A further 47 had a potential ortholog, representing genes with homologous identified by best match BLAST (Pearson, 2013) that were also differentially regulated by cold in Norway spruce but not classified as orthologs by PLAZA (Fig. 1C; Supplemental Table S1). 1395 (72 %) of the Arabidopsis DEGs had identified homologs in Norway spruce that were not cold-responsive in Norway spruce. An additional 167 cold-induced Arabidopsis DEGs had no identifiable homolog in Norway spruce (Arabidopsis singletons). Similarly, of the 3251 cold-induced DEGs in Norway spruce, 401 had orthologs in Arabidopsis and an additional 74 had putative Arabidopsis orthologs. There were 2068 (64 %) Norway spruce cold-induced DEGs with homologs that were not cold-responsive in Arabidopsis and another 708 that had no identifiable homolog in Arabidopsis (Norway spruce singletons). These data revealed a core cold-induced regulon of 387 orthologous genes in Arabidopsis and 475 orthologous COR genes in Norway spruce, which represented 20% and 15% of the cold-induced DEGs, respectively. The orthologous genes were associated with the Gene Ontology (GO) categories “stress response”, “ion binding” and “nucleic acid binding transcription factor activity”, consistent with the known importance of these processes in cold acclimation and demonstrating a conservation of the transcriptional response between these two divergent species (Supplemental Table S2 and S3).

**Figure 1.**
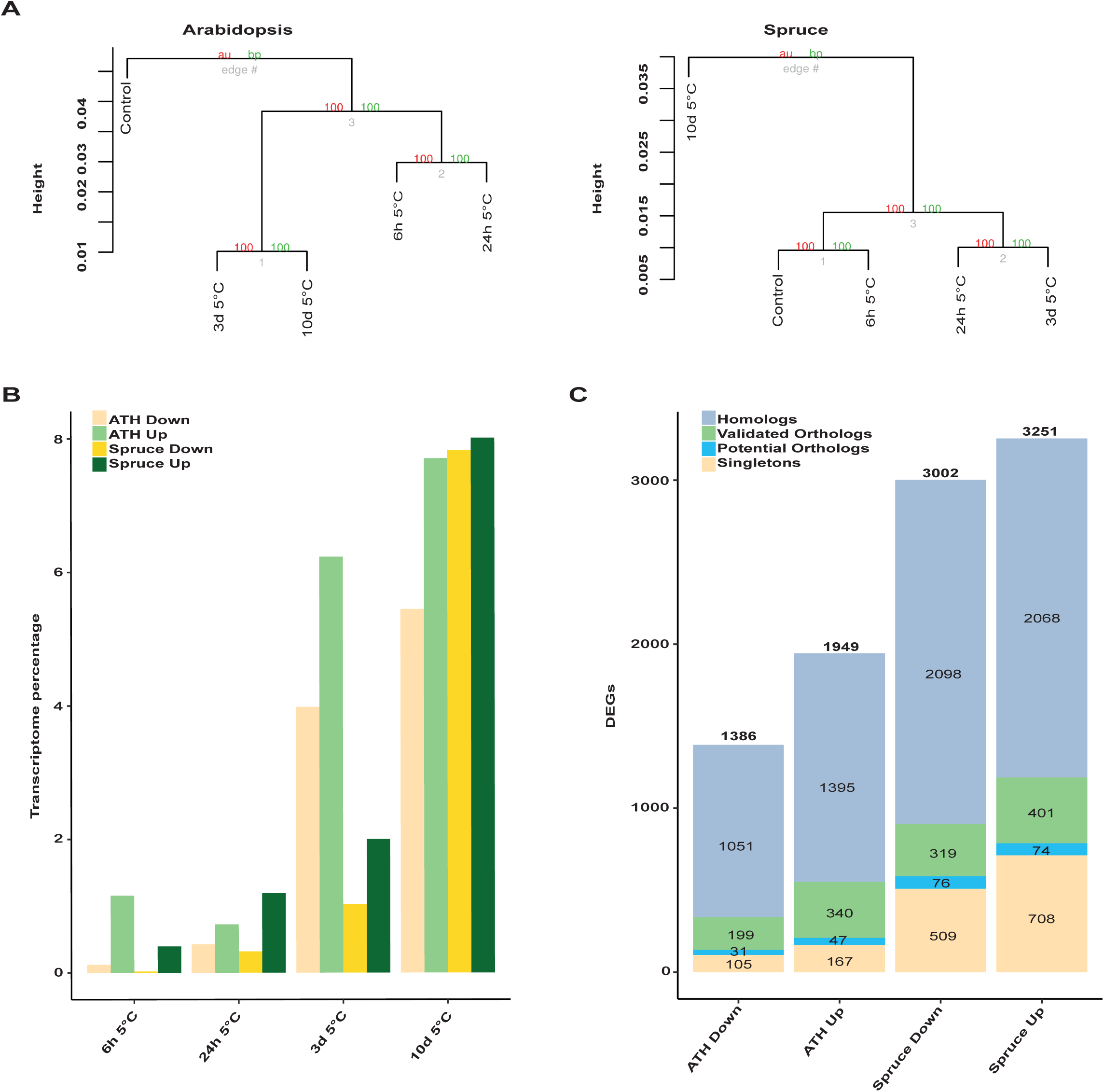
Comparing the response of *Arabidopsis thaliana* leaf and *Picea abies* (Norway spruce) needle transcriptomes exposed to 5°C. **A)** Hierarchical clustering using normalized data (see methods). The red numbers correspond to Approximately Unbiased (AU) values and the green ones to Bootstrap Probability (BP) values. **B)** Analysis of transcriptome progression in response to cold. Differentially expressed gene lists (DEGs) were obtained at each point in the time series, compared against the control, and then represented as a percentage of the transcriptome. DEGs significantly induced in Arabidopsis (green) and Spruce (dark green) and significantly repressed DEGs in Arabidopsis (yellow) and Spruce (dark yellow) were obtained by filtering the data by corrected Pvalue ≤0.01 and Fold Change ≥2. **C)** Orthologs, Homologs and Species-specific DEGs for both species (down and up-regulated). Validated orthologs correspond to orthologous genes that are differentially regulated by cold in both species. Gene lists for each group and functional information are available in Supplemental Table S1.

To compare the functional response of the two species in more detail, we performed DEG enrichment analysis using GOSlim categories. We found that the number of enriched categories increased over time in both species, indicating an extensive remodulation of the transcriptome with the progression of cold exposure (Fig. 2; Supplemental Table S4 and S5). Both species showed significant enrichment of genes involved in “response to stress”, “nucleic acid binding transcription factor activity”, “plasma membrane” and “carbohydrate metabolic processes”. A distinct response in Arabidopsis resulted in significant enrichment of up-regulated genes in the “lipid metabolic process”, “secondary metabolic process”, “cell wall organization or biogenesis” and “catabolic process” that were not significantly enriched in Norway spruce. On the other hand, only Norway spruce showed significant enrichment for “vacuole”, “transport”, “transmembrane transporter activity”, “ion binding”, “cellular protein modification process”, “signal transduction” and “Golgi apparatus” and these processes became significant only after 10 days. Genes within these categories were also induced in Arabidopsis but did not become significant at any time (Fig. 2; Supplemental Table S4 and S5). Among down-regulated genes, Arabidopsis showed strong enrichment for genes related to “structural constituent of ribosome”, “translation” and “RNA binding” that were all largely absent in the Norway spruce, suggesting a strong reorganization of the translational machinery in the herbaceous leaf that was not present in the Norway spruce needle. Similarly, Arabidopsis strongly and rapidly down regulated “plastid” and “photosynthesis” genes but Norway spruce did so much less and much later.

**Figure 2.**
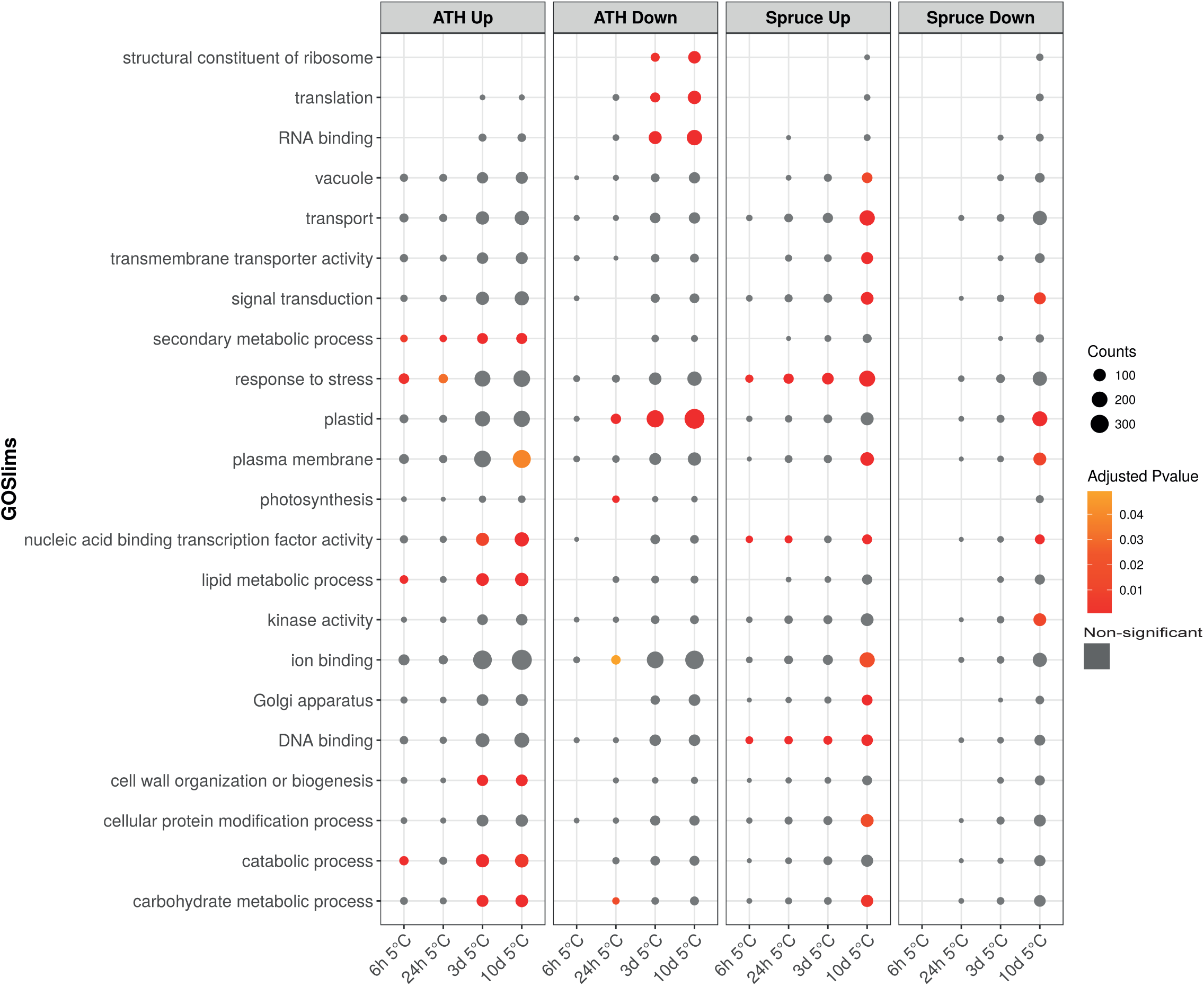
GOSlim functional analysis of the *Arabidopsis thaliana* leaf and *Picea abies* (Norway spruce) needle transcriptomes exposed to 5°C. The number of genes in each category are represented by bubble sizes (counts). GOSlim enrichments with Bonferroni corrected P-values ≤0.05 are represented with a red color scale. Non-significant P-values (corrected P-values >0.05) are in grey. A full list of GOSlim categories and P-values is included in Supplemental Table S4 and a description of the genes and their homologs in Supplemental Table S5.

In order to identify the transcription factors (TFs) that drove the transcriptional responses in both species, the composition and expression profiles of TFs Differentially <u>E</u>xpressed in response to <u>C</u>old (TF-DEC) were analyzed (Fig. 3). First, transcriptional regulators known to respond to cold in Arabidopsis, such as CBFs, were analyzed and showed an induction at 6 h with a fast decline observed by 24 h (Fig. 3A), as shown previously (Chinnusamy et al., 2003; Vogel et al., 2005; Kim et al., 2013; Shen et al., 2015; Kim et al., 2017). Overall, all of the 120 up-regulated TFs in Arabidopsis responded early to cold between 6 and 24 h at 5 °C (Fig. 3B) and most belong to ERF, bHLH and MYB families, which were previously reported as families involved in plant cold and dehydration responses (Fowler and Thomashow, 2002; Chinnusamy et al., 2003; Vogel et al., 2005; Yamaguchi-Shinozaki and Shinozaki, 2006). These Arabidopsis TF-DEC fell into two main clusters; one, including *ERF6* and *MYB7*, with a transient expression peak at 6 h (the top cluster in Fig 3B), and a second larger cluster with a broader expression peak in which expression was induced by 6 h and remained high until 24 h after exposure to cold (bottom cluster, Fig3B). The second cluster included several AP2/ERF superfamily members, such as *TINY2, DEAR2, DEAR4, ERF3* and *AP2* (Fig 3B). This contrasted sharply with what was observed for the 107 up-regulated TF-DEC from Norway spruce, where only a few TFs were induced early (at 6 or 24 h) and these were six *ORA47* homologs, one *AP2*, one *MYB33*, one *ERF53*, one *NAC025* and one *TINY2*-like homolog. The main response of TFs in Norway spruce was delayed relative to Arabidopsis and formed two sub-clusters, one showing induction between 24 h and 3 d and a second sub-cluster with an even later expression maximum at 10 d that was completely absent in the Arabidopsis response. This delayed response in Norway spruce was associated with TFs that belong to AP2/ERF superfamily such as *ERF1, ERF2, ERF9* and *TEM1* homologs, along with other TFs such as *ZFP4; MYB123* and *MYB101*; *anac028* and *anac078* and *WRKY20*-like genes whose role in cold stress response has not been directly reported in Arabidopsis (Fig. 3B). Thus, the results indicated that while Norway spruce shared an early transcriptional response with Arabidopsis at 6 to 24 h, the more extremophile conifer mounted a delayed but more extensive and diverse transcriptional response following 3 and 10 d of exposure to cold.

**Figure 3.**
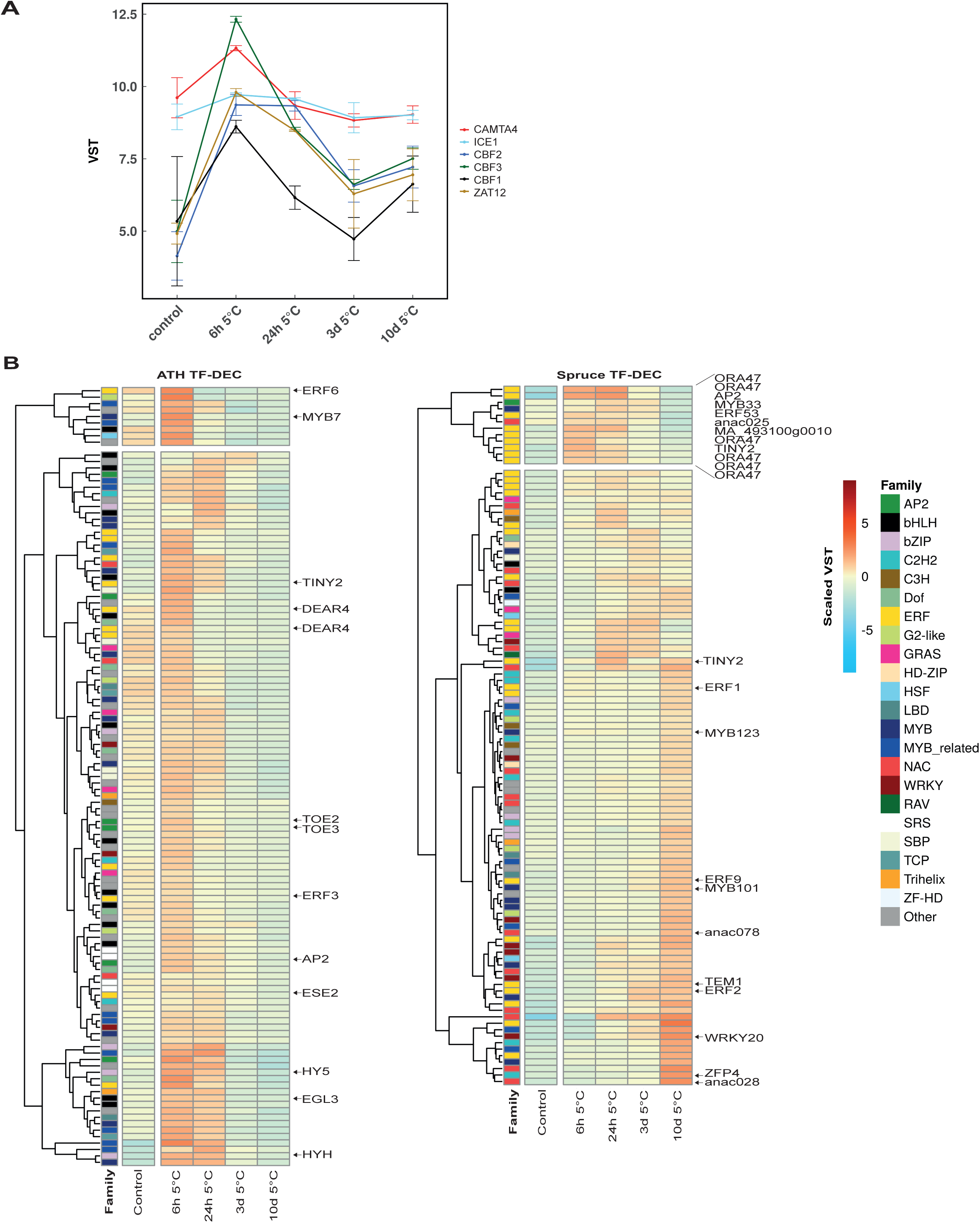
Transcription factor analysis. **A)** Expression profiles of previously characterized Transcription factors (TF) involved in the cold response in *Arabidopsis thaliana: CAMTA4* (AT1G67310), *ICE1* (AT3G26744), *CBF1* (AT4G25490), *CBF2 (*AT4G25470), *CBF3* (AT4G25480) and *ZAT12* (AT5G59820) are shown using variance stabilizing transformation (VST) gene expression values from our experiment (n=3). Errors bars represent SD. **B)** TF differentially expressed by cold (TF-DEC) were analyzed in both Arabidopsis and Norway spruce. TF with positive changes relative to control are shown (corrected P-value ≤0.01 and Fold Change ≥2). VST data were scaled by row means. For each heatmap zoom versions including all the identifiers are available in Supplemental Fig. S6 and S7.

### Analyzing Norway spruce cold stress responses in needles and roots

When the response of Norway spruce needles and roots was compared during exposure to cold (5 °C) and freezing (−5 °C) temperatures, the strongest transcriptional response occurred in both tissues between 3 and 10 d at 5 °C (Fig. 4A). Differential expression analysis revealed a progressive increase in the number of up-regulated DEGs in both tissues during the first acclimation phase up to 10 d at 5 °C (Fig. 4B). Although the root response lagged behind at 6 h, presumably due to the effect of soil acting as a temperature buffer, after 24 h the numbers of DEGs were largely similar between the two tissues and the largest response in both tissues occurred between 3 and 10 d in the cold. Subsequent exposure of both tissues to freezing at −5 °C had no noticeable effect on the number of DEGs in needles but did result in a further increase in root DEGs after 3 and 10 d at −5 °C, giving a total of 4324 and 4407 up-regulated DEGs in needles and roots respectively (Fig. 4B).

**Figure 4.**
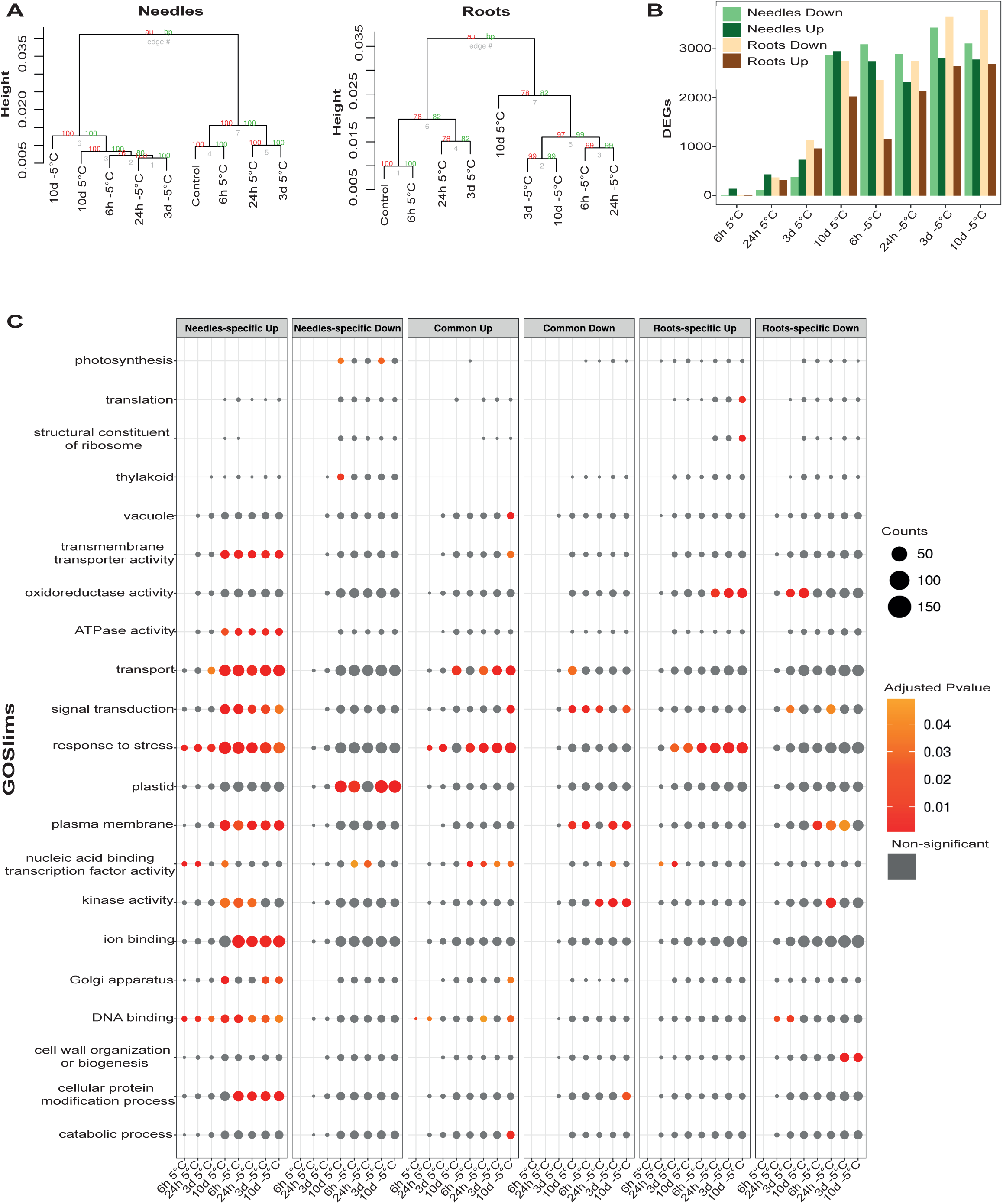
The response of needles and root transcriptomes from Norway spruce to exposure to cold (5°C) and freezing (−5°C). **A)** Hierarchical clustering of gene expression data from needles and roots samples (see methods). Red numbers correspond to Approximately Unbiased (AU) values and green ones to Bootstrap Probability (BP) values. **B)** Analysis of transcriptome progression in response to cold (5°C) and freezing (−5°C). Differentially expressed gene lists (DEGs) were obtained at each point in the time series, compared against the control. DEGs in Needles (dark green represents induced genes and light green represents repressed genes) and Roots (brown represents induced genes and tan represents repressed genes) were obtained by filtering the data by corrected Pvalue ≤0.01 and Fold Change ≥2. **C)** The number of DEGs (counts) with different GOSlim tags assigned are represented by bubble size. GOSlim enrichments with Bonferroni corrected P-values ≤0.05 are represented with a red color scale. Non-significant P-values (corrected P-values >0.05) are in grey. A full list of GOSlim enrichments is included in Supplemental Table S6 and a complete description of each GOSlim category and the gene functions in Supplemental Table S7.

GOSlim enrichment analysis showed that while both needles and roots deployed similar numbers of up-regulated DEGs and most GOSlim categories were common between the tissues, in needles more categories were significantly enriched. This lack of common enrichment demonstrates differential tissue-specific responses by needles and roots to cold (Fig. 4C). Even in the “response to stress” category, while there was significant enrichment in up-regulated common DEGs, there were stronger tissue-specific responses, especially in needles (Fig. 4C). Similarly, genes in the “transport” category were enriched in up-regulated DEGs common to both tissues and also in needle-specific genes, showing that the transport of solutes and ions across membranes is important in both tissues, but more marked for needle-specific response genes. This stronger response by needles is further demonstrated by the finding that “plasma membrane”, “transmembrane transporter activity” and “ion binding” categories were only enriched in the up-regulated needle-specific DEGs.

Unsurprisingly, in the down-regulated DEGs photosynthesis and plastid categories were the main gene classes significantly enriched in the needle specific list. In addition, downregulation of “nucleic acid binding transcription factor activity” genes at 6 and 24 h under freezing, reflected deactivation of bHLH, bZIP, MYB, NAC and AP2/ERF superfamily TFs in needles (Fig. 4C; Supplemental Table S6 and S7). On the other hand, root-specific and common down-regulated DEGs were enrichment in “plasma membrane”, “kinase activity” and “signal transduction” categories, indicating that the silencing of some components of these categories occurred in both tissues.

### Characterization of the induced response to cold in Norway spruce

We identified 4324 up-regulated DEGs in needles and 4407 in roots of which 2002 were common between both tissues (Fig. 5A). To identify genes with similar transcriptional responses, we performed hierarchical cluster analysis on these up-regulated DEGs and identified the main clusters in needle-specific, common and root-specific DEGs, identifying 7, 11 and 8 main clusters, respectively (Fig. 5B). When we applied correlation and cluster analysis to all identified clusters, 7 main responses were found, termed <u>S</u>uper <u>C</u>lusters (SC) (Fig. 5C and Supplemental Table S8). SC-6 contained Cluster N1 after best correlation with itself only. Its early response showed a needle-specific behavior characterized by an early induction, followed by expression falling below control levels. Similarly, SC-7 was an early response cluster, formed by Cluster N2, Cluster R4 and Cluster C6 (as expressed in needles). These two SCs (SC6 and SC7) represented early coordinated responses to cold. Expression of genes in SC-1, SC-2 and SC-3 steadily increased during exposure to 5 °C, and represented genes likely responding downstream of SC6 & 7; SC-2 was an exception and contained differentially expressed genes in roots only. SC-4 corresponded to Cluster R3 and the behavior of this SC showed a progressive increase in gene expression until 10 d 5 °C and then a continuous drop during freezing. SC-5 showed a similar response pattern to SC4 but was characterized as a common response in both tissues defined in Cluster C9. In general, there were early and late response clusters in both tissues, but with maximum induction levels at different time points, supporting the conclusion that distinct needle and root responses exist.

**Figure 5.**
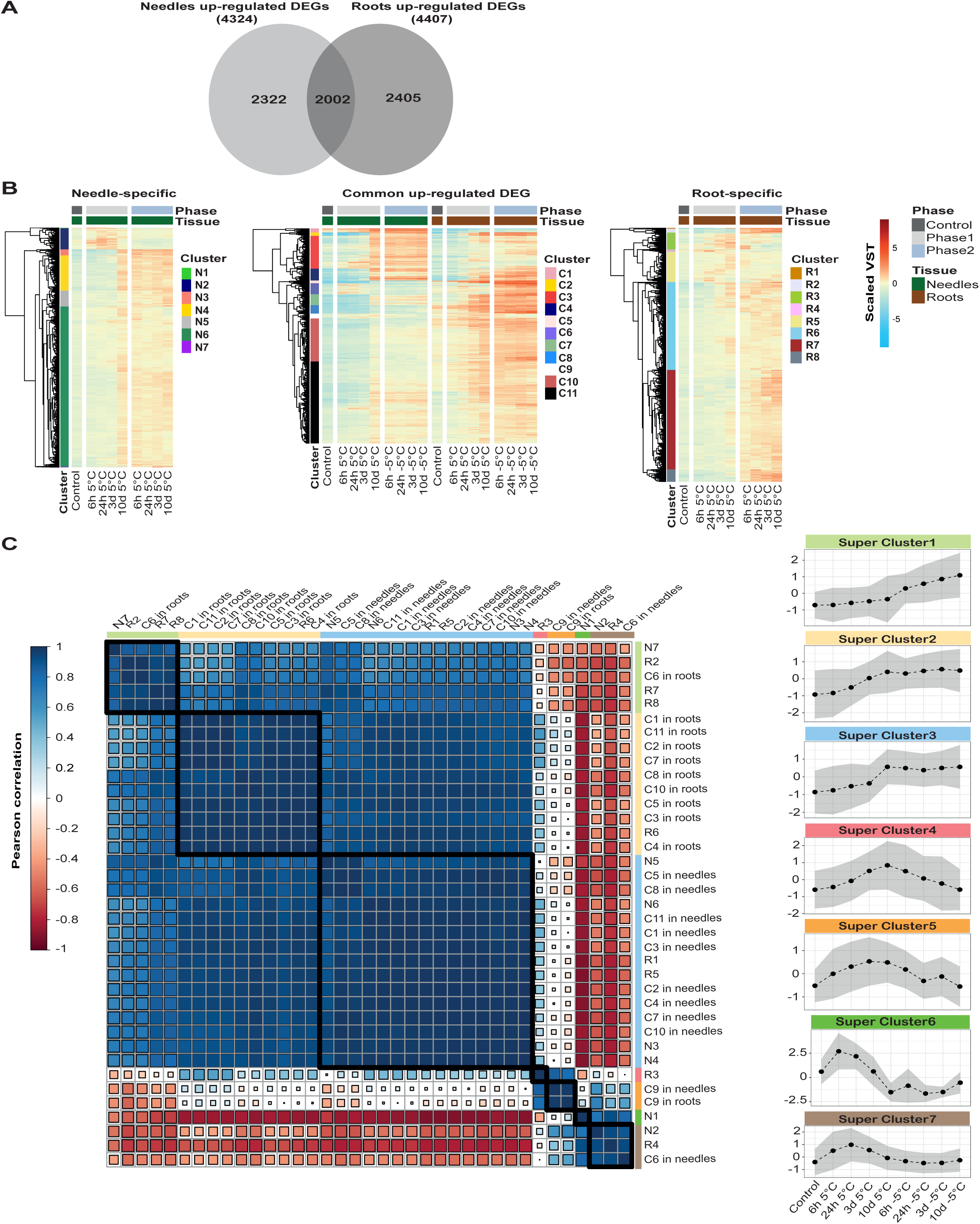
Identification of coordinated responses to cold. **A)** Venn diagram representing needle-specific, root-specific and common up-regulated differentially expressed genes (DEGs). A GO enrichment analysis of these gene lists is in Supplemental Table S7. **B)** Heat maps and main clusters of tissue specific and common up-regulated DEGs. **C)** Super clusters (SC) were defined by combining clusters from Fig 5B using *Pearson* correlation analysis. Common clusters were separated by tissues to compare against tissue-specific clusters. Scaled gene expression of each SC and their mean gene expression is represented with dotted lines. Grey areas represent variability by two standard deviations. An analysis of SC distribution in the network is available in Supplemental Figure S3. A Gene Ontology (GO) enrichment analysis for these super clusters is available in Supplemental Table S8.

The transcription factors associated with the cold responses in Norway spruce needles and roots were examined and 215 TFs induced in response to <u>c</u>old (TF-DEC) were identified from needles and 183 from roots, of which 107 (37%) were needle-specific, 75 (26%) root-specific and 108 (37%) common to both tissues. In addition, 20% (57 genes) of these were Norway spruce-specific (singletons). These included 15 TFs without any family assigned, three ERF, one bZIP and one WRKY and also TFs families that have not previously been implicated in stress tolerance, such as 15 HD-ZIP, six GRAS, four TALE and three C2H2, among others (Baillo et al., 2019) (Supplemental Table S9). However, while these singletons all responded to cold stress, none showed strong induction or repression in response to either cold or freezing (Supplemental Fig. S4) and from this analysis there is no evidence to suggest these singletons represent novel transcription factors that contribute to the enhanced tolerance of evergreen conifers. Clustering of the expression profiles of these TFs resolved two main clusters in the needle-specific TF-DEC (Fig. 6); cluster *I* was comprised of TFs induced 3 to 10 days after exposure to cold; and a second smaller cluster *II* was induced earlier between 6 h and 3 d and these TFs returned to pre-cold exposure expression levels or lower by day 10. No freezing-specific induced TFs were found in the needle-specific group. In contrast, the root-specific TF-DECs clustered into four primary groups; cluster *III* showed progressive induction during cold stress and this was largely maintained during subsequent freezing, cluster *IV* included TFs induced between 24 h and 3 d and lower expression levels following freezing; clusters *V* and *VI* which, unique to roots, contained TFs that were strongly induced following exposure to freezing at −5 °C (Fig. 6 and Supplemental Table S10).

**Figure 6.**
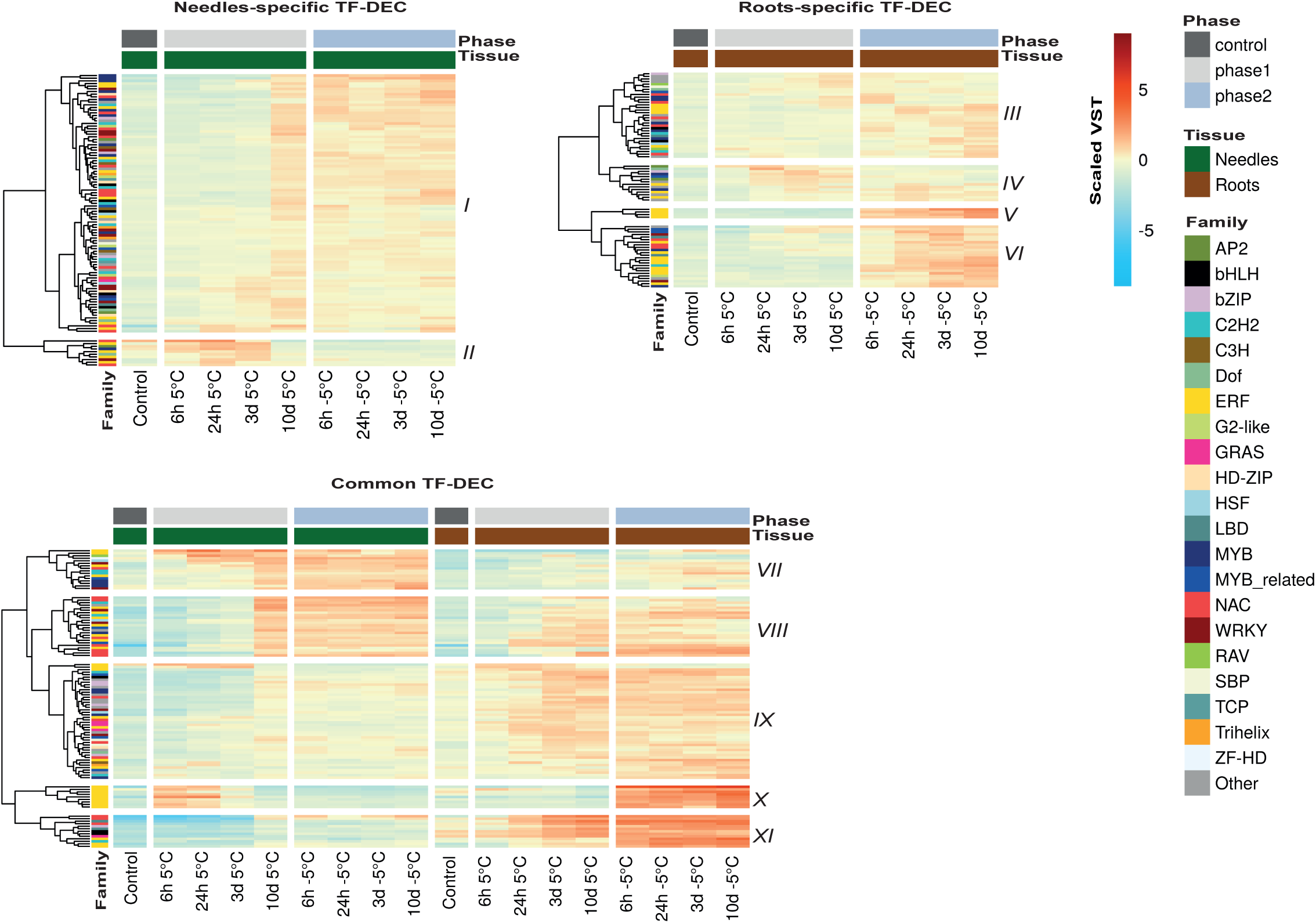
Transcription factors differentially regulated by cold (TF-DEC). TF-DEC were analyzed by heat maps and hierarchical clustering (corrected P-value ≤0.01 and Fold Change ≥2). Family members were obtained from the Plant Transcription Factor Database (Jin, Zhang et al. 2014) and normalized VST expression values were scaled by row means. A file including gene descriptions for each cluster is available in Supplemental Table S10.

Within the common TF-DECs, five primary groups were identified; cluster *VII* included TFs induced early in needles and that maintained high expression levels during exposure to freezing. In contrast, these TFs were only weakly induced in roots and mostly during exposure to freezing; cluster *VIII* included TFs not induced in needles until 10 d in the cold and thereafter expression levels remained high during freezing, while in roots these TFs are induced starting from 24 h exposure to cold; cluster *IX* included TFs only weakly responsive in needles but these TFs showed stronger induction in roots and this was maintained following exposure to freezing; cluster *X* was comprised of a group of ERF TFs that were induced strongly in needles at 6-24 h in the cold and then returned to control levels but were only induced in roots after freezing; lastly cluster *XI* TFs were induced in both tissues but appeared to play a strong role only in roots. Thus, even though there were a large number of common TFs induced in both tissues by exposure to cold and freezing, the majority of these showed strong differential regulation between the two tissues, either in their timing or the strength of their induction.

### Cold response regulatory network of Norway spruce

In order to analyze in detail how cold-responsive (COR) genes were regulated and to identify TF “hubs” playing a key role in COR gene regulation, we performed a regulatory network analysis of TF-DEC and COR genes combining co-expression and promoter motif analysis of all COR genes. A network model of inferred transcriptional regulations of COR genes was built using an approach previously used to identify putative plant cold acclimation pathways and key TFs in Arabidopsis and rice (Chawade et al., 2007; Lindlof et al., 2009). The analyzed motifs were consensus target sequences for ERF, NAC, MYB, bZIP, bHLH, AP2, DRE, LBD and WRKY families, which have previously been associated with plant cold stress (Chawade et al., 2007; Peng et al., 2015). For each motif, confidence intervals (CI) were obtained (Table 1) and used as a criterion of over-representation. Thus, regulatory interaction between two nodes (genes) was the result of a combination between TF-target co-expression and the over-representation of the recognized motif for the respective TF in the target COR gene promoter. Using a strict correlation threshold, the resulting regulatory network included 2135 links of TF and target genes and 910 nodes (genes). 784 were COR target genes and 126 were TFs, with mainly bHLH, MYB, NAC, and ERF family members (Fig. 7A). Analysis of the connectivity of the nodes (degree) in the network showed that the network followed a scale free behavior (Fig. 7B), with only few genes highly connected. These node degree measurements were then used as a gene prioritization criterion (Moreau and Tranchevent, 2012) and the top ten most connected nodes (hubs) were analyzed, all of which were TFs (Fig. 7C). The most connected hub corresponded to the gene MA_448849g0010; a bHLH TF (*ICE1*-like) that putatively regulated (on the basis of TF-COR coexpression and TF binding motif presence) 159 COR genes. Interestingly, the third most connected hub (MA_68586g0010) was a *bHLH101*-like that was only differentially regulated in roots and 88% of the target genes were also root-specific (Fig. 7C and Supplemental Table S13). These results demonstrated that the cold regulatory network in Norway spruce was highly interconnected, and most of the cold response circuits were common to both tissues, including homologs of genes previously reported as regulators of stress response genes in other species such as Arabidopsis (Supplemental Table S13 and Supplemental Fig. S5). A notable exception to this was the root hub *bHLH101* homolog (MA_68586g0010), which suggested that tissue-specific responses are present in this species although the function of this hub and its downstream genes remain to be elucidated.

**Figure 7.**
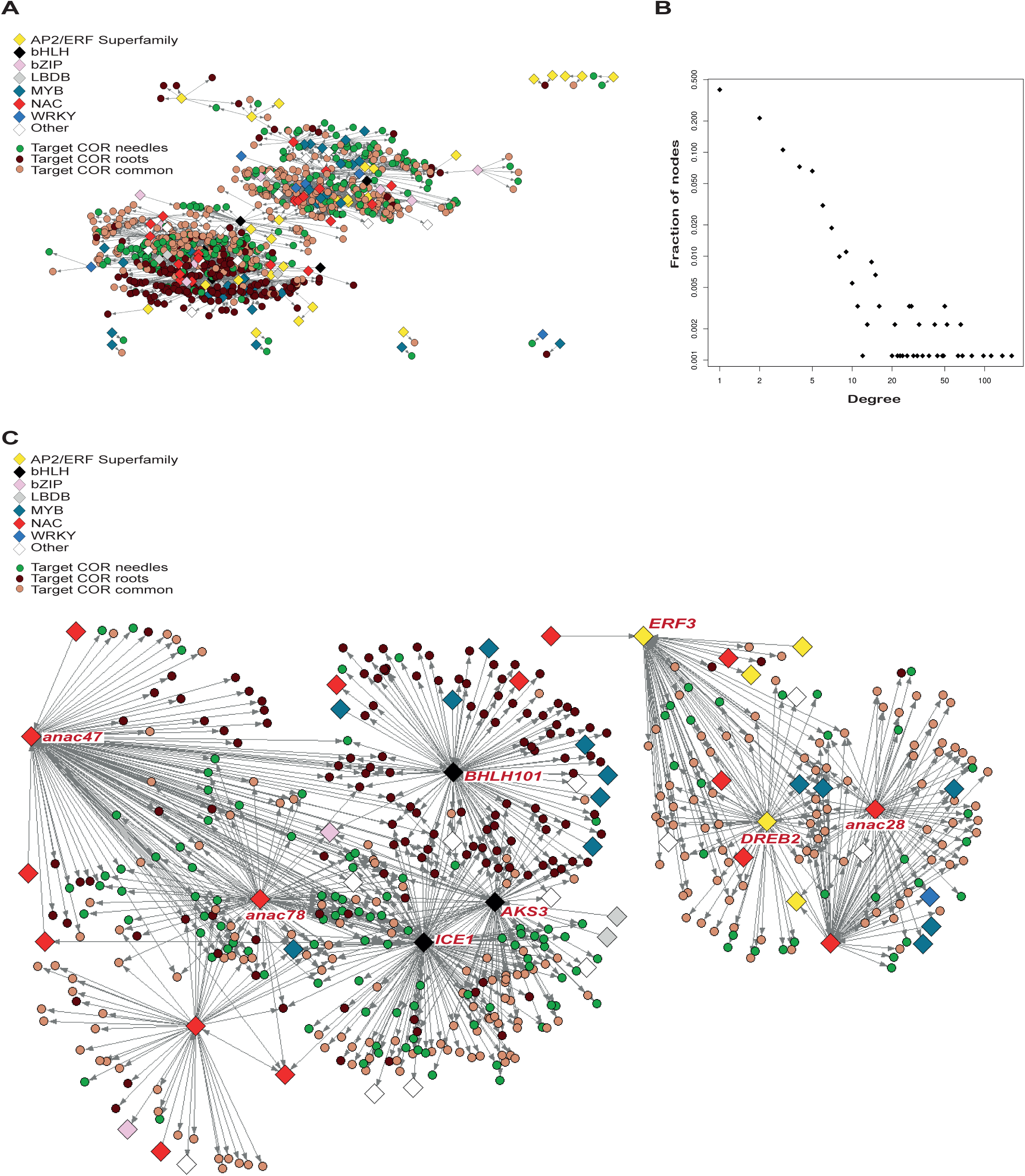
Regulatory network analysis. **A)** Network representation of predicted regulatory interactions between Transcription Factors (TF) and cold responsive (COR) genes. TFs are represented by diamonds and their family by colors. COR genes are represented by circles and colored according to the tissue in which they are differentially regulated. **B)** Network Degree distribution in Log10/Log10 scale. **C)** Sub-network of the 10 genes with the highest centrality. Gene Ontology (GO) enrichments in the hub neighborhoods are available in Supplemental Table S12 and topology information and gene aliases are available in Supplemental Table S13.

**Table 1.**
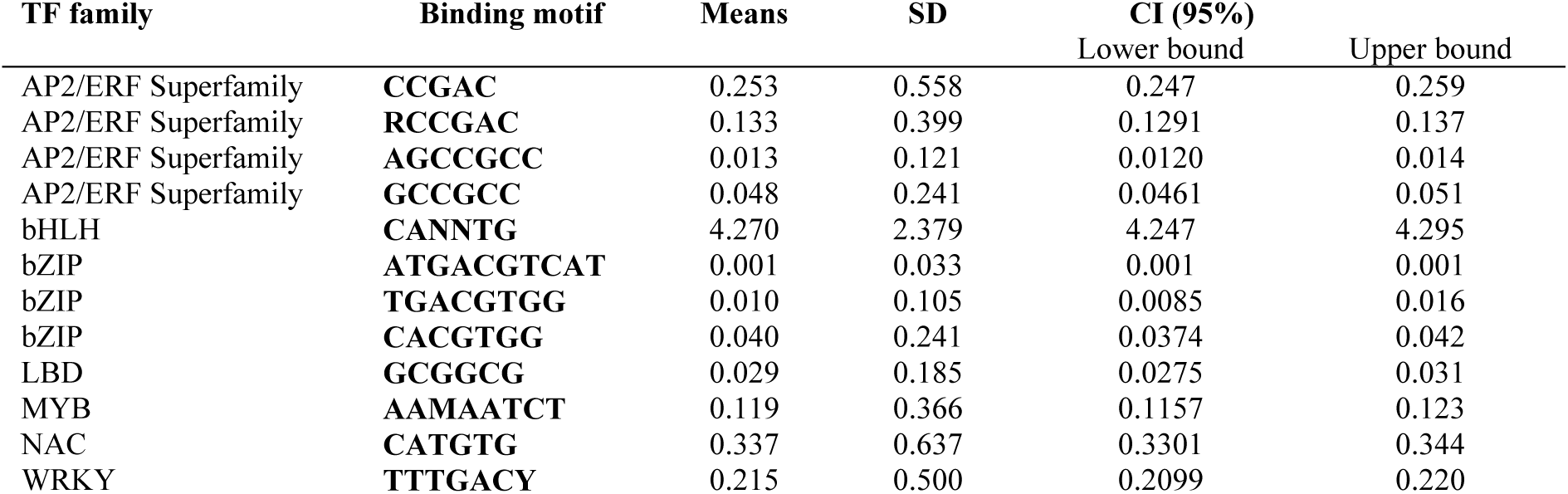
Analyzed transcription factors and consensus binding site motifs. TF selected for regulatory network analysis are presented here. Binding motif were analyzed according with IUPAC nomenclature (Cornish-Bowden, 1985); ‘M’=A or C, ‘N’= any nucleotide A or T or C or G, Y=C or T; CI= Confidence interval.

| TF family | Binding motif | Means | SD | CI (95%) |  |
| --- | --- | --- | --- | --- | --- |
|  |  |  |  | Lower bound | Upper bound |
| AP2/ERF Superfamily | **CCGAC** | 0.253 | 0.558 | 0.247 | 0.259 |
| AP2/ERF Superfamily | **RCCGAC** | 0.133 | 0.399 | 0.1291 | 0.137 |
| AP2/ERF Superfamily | **AGCCGCC** | 0.013 | 0.121 | 0.0120 | 0.014 |
| AP2/ERF Superfamily | **GCCGCC** | 0.048 | 0.241 | 0.0461 | 0.051 |
| bHLH | **CANNTG** | 4.270 | 2.379 | 4.247 | 4.295 |
| bZIP | **ATGACGTCAT** | 0.001 | 0.033 | 0.001 | 0.001 |
| bZIP | **TGACGTGG** | 0.010 | 0.105 | 0.0085 | 0.016 |
| bZIP | **CACGTGG** | 0.040 | 0.241 | 0.0374 | 0.042 |
| LBD | **GCGGCG** | 0.029 | 0.185 | 0.0275 | 0.031 |
| MYB | **AAMAATCT** | 0.119 | 0.366 | 0.1157 | 0.123 |
| NAC | **CATGTG** | 0.337 | 0.637 | 0.3301 | 0.344 |
| WRKY | **TTTGACY** | 0.215 | 0.500 | 0.2099 | 0.220 |

## Discussion

A qualitative comparative analysis of Norway spruce needles with leaves of the temperate herbaceous model plant Arabidopsis showed that both species share a core of orthologous genes that respond to cold (Fig. 1). The orthologous responses include the up-regulation of several TFs of the ERF3 family, including TCP2 and HB13, that are known to play a role in plant stress responses (Stockinger et al., 1997; Liu et al., 1998; Nakano et al., 2006; Chawade et al., 2007; Cabello et al., 2012) (Supplemental Table S3). In addition, construction of a gene regulatory network demonstrated that an ICE1-like TF was, like its counterpart in Arabidopsis, a central player in mediating the cold response in needles and roots of Norway spruce. However, Norway spruce also demonstrated a strongly delayed transcriptional response relative to Arabidopsis, and this delayed response was associated with the induction of a large cohort of TFs, including *ERF1*, *ERF3*, *anac028* and *anac078* homologs, that have not previously been described to have a function in cold acclimation. In addition, a *bHLH101-like* TF was shown to be co-expressed with a root-specific subset of genes in the gene-regulatory network. The Arabidopsis ortholog of this *bHLH101-like* TF was reported to be involved in iron homeostasis (Wang et al., 2007) and photo-oxidative stress responses (Noshi et al., 2018). A function for this gene as a central regulator of cold acclimation has not been described but the strong oxidative-stress response shown by roots (Supplemental Table S11) indicates that this TF may play such a role in coordinating this response in spruce roots, which is supported by the fact that its first degree neighborhood is enriched in oxidoreductase activity genes (Supplemental Table S12). Furthermore, no central regulators have previously been identified as root specific and the findings presented here indicate that tissue specific responses are important in the cold hardiness response, at least in perennial species such as conifers (Figs. 4, 5 & 6 and Supplemental Fig. S10).

Evergreen plants such as Norway spruce maintain their needles above the snow pack during winter and thus require mechanisms to protect the needles from extreme low temperatures and the associated desiccation. On the other hand, roots face less extreme temperatures (Supplemental Fig. S6) but they must also be protected from freezing damage. To date, metabolomic and proteomic analyses have shown that, as for herbaceous leaves, carbohydrate and lipid metabolism and the accumulation of dehydrins play key roles in survival at extreme low temperatures in perennial evergreen needles (Kjellsen et al., 2013; Angelcheva et al., 2014; Strimbeck et al., 2015). In our analysis, we found that both tissues mounted a progressive response to cold and that following subsequent freezing, most common cold acclimation responses were maintained in both tissues. GO enrichment analysis performed on all DEGs that are induced in Norway spruce revealed that regardless of the tissue, common genes responding positively to cold were over-represented for genes within the core stress categories of “regulation of transcription”, “transport” and “response to wounding” (Fig. 5A; Supplemental Table S11). However, some responses were greater in needles or roots or became more important in response to freezing (Fig. 4C). GO enrichment analysis showed that needle-specific induced genes were enriched for “plasma membrane”, “ATP-binding”, “DNA binding”, “integral component of membrane” and “Golgi apparatus”. On the other hand, root-specific induced DEGs were enriched for the categories “peroxidase activity”, “oxidation-reduction process”, “metal ion binding” and “structural constituent of ribosome” and “translation” (Supplemental Table S11), consistent with earlier findings that roots are prone to hypoxia during winter due to ice-encasement (Martz et al., 2016). These contrasting responses indicated that while needles are exposed to extreme cold and desiccation, demanding membrane reorganization and protection; roots, which remain more metabolically active during winter (Law BE, 1999; Martz et al., 2016), are under higher oxidative stress.

The best studied of the cold response pathways in herbaceous species is that regulated by the CBF/DREB1 (CRT-binding factor/DRE-binding protein) family of TFs, which belong to the ERF family (Shinozaki and Yamaguchi-Shinozaki, 1996; Fowler and Thomashow, 2002; Cook et al., 2004; Jin et al., 2014). In Arabidopsis the CBFs show a transient early induction by cold in roots and leaves (Kilian et al., 2007; Hruz et al., 2008). A single ortholog of CBF1 and CBF3 was found in Norway spruce (MA_20987g0010), which was induced between 6 and 24 h at 5°C in roots, although it did not pass the statistical filters to be classified as an up-regulated DEG or COR gene and it was not responsive to cold in needles (Supplemental Fig. S7). Cluster X of the common TF-DEC (Fig. 6) contained nine ERF TFs showing early induction in needles, indicating that other genes of the same TF family have a role in cold stress regulation in aerial tissues of Norway spruce, and that this stress response may be supported by an expansion of the ERF family of TFs in Norway spruce (Supplemental Fig. S8 and S9). Based on co-expression and *cis*-element over-representation analysis, we built a regulatory network to identify potential key conserved TFs mediating cold acclimation in Norway spruce (Fig. 7). We found extensive crosstalk between the COR genes, with TFs being regulated by upstream TFs and also a module with a root-specific bias. Interestingly, the most connected TF in the network was an *ICE1*-like gene (MA_448849g0010), orthologous to an upstream bHLH transcriptional activator of CBFs genes in Arabidopsis that is activated by the kinase OST1 under cold stress (Chinnusamy et al., 2003; Ding et al., 2015). Our results indicate that this Norway spruce *ICE1*-like protein is a potential regulator of many COR genes, 150 of which are genes expressed in both analyzed tissues (Supplemental Table S13), and its progressive late induction suggests that its role may become even stronger following freezing (Supplemental Fig. S10).

In conclusion, our study provides a comprehensive overview of the transcriptome of Norway spruce involved in cold stress and reveals new mechanisms to face extreme low temperatures, which are especially important to explain how Norway spruce is able to overcome extreme winters. We additionally provide new candidate genes to inform the design and interpretation of future studies, such as target COR genes and TFs. Overall, the identified TF-DEC and hubs obtained from our regulatory network can be used to identify the most suitable candidate genes for carrying out genetic modifications or directed breeding to generate high-yielding cold stress tolerant trees, which potentially could solve many problems in the forest industry in the face of the new expected global change scenarios.

## Material and methods

### Experimental design and sampling

Two-year-old Norway spruce (*Picea abies* (L.) Karst.) seedlings of the seed provenance Lilla Istad (56° 30’ N) were planted in peat in 3 l pots and grown at 18 °C/15 °C light/dark in a 16-h light period. Needles and roots of five seedlings were sampled at control conditions. Forty seedlings were shifted to continuous cold temperature at 5 °C for ten days. Needles and roots were collected 6 h, 24 h, 3 d and 10 d after the start of the cold treatment. Subsequently the seedlings were exposed to sub-zero freezing temperature at −5 °C and samples were taken 6 h, 24 h, 3 d and 10 d. Samples were always taken from previously unsampled plants. Every sampling point was represented by needles from five seedlings, three of which were also sampled for roots. Samples were collected on dry ice and stored at −80 °C until further processing.

Seeds of *Arabidopsis thaliana* (Col-0) were sown into soil and kept in an 8 h photoperiod (150 μmol photons m^-2^ s^-1^) at 23°C. After 14 days seedlings were transplanted into individual pots. After a further 30 days plants were shifted to 5°C and leaves of these plants were sampled after 0, 6, 24 hours and 3 and 10 days at 5°C. Leaves were harvested from 5-10 randomly chosen plants and were pooled. Three pools were collected per time point.

### RNA preparation and sequencing

Norway spruce samples were prepared by CTAB method (Chang, 1993) with following modifications: Addition of warm extraction buffer, including polyvinyl pyrrolidinone (PVP) 40, and vortex mixing of the ground sample material was followed by an incubation step for 5 min at 65 °C. Precipitation with ¼ volume 10 M LiCl took place at −20 °C for 2 h and RNA was then harvested by centrifugation at 14.000 rpm for 20 min and 4 °C. The RNA was further purified using the RNeasy mini kit (QIAGEN, Hilden, Germany). A DNase Digestion with the RNase-free DNase set (QIAGEN) was included in the procedure. High quality total RNA with a RIN ≥ 7.5, OD 260/280 ratio of ≥ 2.0 and concentrations ≥ 50 ng/µl was sequenced by SciLifeLab (Stockholm, Sweden) for paired-end (2 x 125 bps). The sequencing library preparation included an enrichment for poly-adenylated mRNAs and all samples yielded > 9.4 million read pairs.

*Arabidopsis thaliana* (Col-0) leaf samples were isolated using the Plant RNA Mini kit (E.Z.N.A.) according the manufacturer’s instructions. DNase treatment was performed after RNA extraction using DNA-free^TM^ DNA Removal Kit (Ambion, Life Technologies). Samples were sequenced by BGI (Beijing Genomics Institute) and all yielded > 23 million paired-end reads.

All RNA samples integrity were analyzed by Agilent RNA 6000 Nano kit (Agilent Technologies, Waldbronn, Germany) on a Bioanalyzer 2100 (Agilent Technologies) and purity measured with a NanoDrop 2000 spectrophotometer (Nanodrop Technologies, Wilmington, DE, USA). All samples passed quality controls are were sequenced by Illumina HiSeq 2000 platform.

### Data processing

The reads pre-processing and quality assessment of the raw data was performed as in Delhomme et. al. (2014) following the standard guidelines http://www.epigenesys.eu/en/protocols/bio-informatics/1283-guidelines-for-rna-seq-data-analysis <u>using FastQC-0.10.1 https://www.bioinformatics.babraham.ac.uk/projects/fastqc/ for quality control and SortmeRNA-2.0 to filter RNA contaminants (Kopylova et al., 2012). Trimmomatic-0.022 (Bolger et al., 2014) was used for trimming and adapter removal and STAR-2.4.0f1 for read alignments (Dobin et al., 2013). To summarize read counts per transcript and obtain count data we used HTSeq-0.6.1 (Anders et al., 2015) and to identify candidate genes that were differentially expressed and to obtain variance stabilizing transformation (VST) normalized expression values we used DESeq2-1.16.1 package (Love et al., 2014). Hierarchical clustering were performed using Pvclust R package (Suzuki and Shimodaira, 2006).</u>

### Data availability

Sequencing data are available at the European Nucleotide Archive (ENA) as accession PRJEB26934. Code for reproducibility is availability at the Github page https://github.com/avergro/Spruce-cold-stress. The expression profiles of Norway Spruce and Arabidopsis genes can be viewed interactively through a web service at https://hurrylab.shinyapps.io/spruce-cold-stress.

### Functional analysis

The predicted TF genes and its associated families in Norway spruce were obtained from PlantTFDB 3.0 (Jin et al., 2014). Norway spruce sequences and Gene Ontology annotations were obtained from Conifer Genome Integrative Explorer (ConGenIE; http://congenie.org) (Nystedt et al., 2013). Gene Ontology and GOSlim enrichment analyses were performed with standalone version of GeneMerge (Castillo-Davis and Hartl, 2003) using as background population a transcriptome of 43398 genes. To compare globally the functions against *Arabidopsis thaliana* the GOSlim tags were assigned to the Norway spruce genes using the plant slims subset available in Gene Ontology Consortium web site (Ashburner et al., 2000; Beike et al., 2015) by Map2Slim function from Owltools https://github.com/owlcollab/owltools/wiki/Map2Slim. To identify previously characterized COR genes in Norway Spruce, a bidirectional BlastP (Altschul et al., 1990) using *Arabidopsis thaliana* (TAIR 10) (Garcia-Hernandez et al., 2002; Rhee et al., 2003) and *Picea abies* v1.0 references proteomes (Nystedt et al., 2013), which is available in ConGenIE (Sundell et al., 2015) was used. Then, best Blast matches were used to assign gene aliases. Peptides sequences of 10 or less amino acids were removed. A total amount of 58.587 amino acid sequences were analyzed in Norway spruce after excluding low confidence sequences. Orthology analysis was performed using PLAZA resource (Van Bel et al., 2018).

### Regulatory network analysis to find pivotal TF of cold stress response

The upstream regions of Norway spruce gene models containing start and stop codons were obtained from the FTP web site available in ConGenIE and dropped all of them at the same lengths of 1Kb using a Perl script. Genes with upstream regions shorter than 1Kb were excluded of the analysis. In total 37621 upstream regions were analyzed. Counts of the different analyzed motifs in the promoters regions (Table 1) were obtained with the stand-alone version of PatMatch (Yan et al., 2005). A motif was considered over-represented in a target gene promoter if was present in an upstream region more than the respective confidence interval (CI) upper bound of the abundance of the different analyzed motif (Chawade et al., 2007). The analyzed motifs were consensus target sequences for ERF, NAC, MYB, bZIP, bHLH, AP2, DRE, LBD and WRKY families. The upper limit of the confidence intervals was below 1, with an exception for bHLH where the upper limit was over 4. In these cases, just a single occurrence was considered as a case of over-representation because these sequences are not sporadically distributed on the promoter regions. For bHLH five or more times was considered as over-representation of the motif in a promoter. We got the co-expression values between all the identified COR genes and identified TF from Norway spruce using Spearman correlations values performing significance test of the correlation coefficients and FDR corrected P-values were obtained for them. A threshold of Spearman correlation ≥ 0.8 and adjusted P-value ≤ 0.01 was used to filter the correlations. A regulatory interaction in our network was obtained by the combination of a co-expression data and over-representation of binding motifs. So, if a pair TF-Target gene overcome a correlation threshold and the target gene has an over-representation of the motif, which recognize the TF in the promoter, then these TF-Target gene pair make up a regulatory interaction in our regulatory network. Pajek software was used to obtain network parameters and visualizations (Nooy et al., 2005).

## Author contributions

VH and NRS conceived and supervised the project; AV, JCH and PS performed the research; AV conducted the bioinformatics analysis; AV, JCH, NRS and VH wrote the manuscript, which was edited by all authors.

## Acknowledgements

The authors wish to thank the UPSC bioinformatics facility (https://bioinfomatics.upsc.se) for technical support with regards to the RNA-Seq data pre-processing and analyses. We would also like to acknowledge support from Science for Life Laboratory, the Knut and Alice Wallenberg Foundation, the National Genomics Infrastructure funded by the Swedish Research Council, and Uppsala Multidisciplinary Center for Advanced Computational Science for assistance with massively parallel sequencing and access to the UPPMAX computational infrastructure.

## Supplemental Data

The following supplemental information is available:

**Supplemental Figure S1.** Arabidopsis TF-DEC.

**Supplemental Figure S2.** Spruce TF-DEC.

**Supplemental Figure S3.** Super cluster (SC) member distribution in the predicted regulatory network.

**Supplemental Figure S4.** Heatmap of Norway spruce-specific transcription factors differently regulated by cold (TF-DEC).

**Supplemental Figure S5.** Aliases of the first neighborhood genes of the 10 hubs with the highest centrality in the predicted regulatory network.

**Supplemental Figure S6.** Annual air and soil temperature measurements in a typical northern Norway spruce forest..

**Supplemental Figure S7.** Expression profile of a putative CBF ortholog in Norway spruce.

**Supplemental Figure S8.** TF family distribution.

**Supplemental Figure S9.** Distribution of Norway spruce TF-DEC families.

**Supplemental Figure S10.** Heatmaps of expression levels of transcription factors differently regulated by cold (TF-DEC).

**Supplemental Table S1.** Orthologs, potential orthologs and singletons identified between *Picea abies* and *Arabidopsis thaliana*.

**Supplemental Table S2.** GOSlim enrichments analysis on orthologs and potential orthologs gene lists.

**Supplemental Table S3.** Selected functional overlaps between Arabidopsis and Spruce.

**Supplemental Table S4.** GOSlim enrichments in differential expressed gene lists of Spruce and Arabidopsis over time series treatments.

**Supplemental Table S5.** GOSlim terms and gene lists in vertical format of Spruce and Arabidopsis over time series treatments including gene descriptions.

**Supplemental Table S6.** GOSlim enrichments in differential expressed gene lists (DEGs) of Spruce over time series treatments analyzing needle-specific, root-specific and common DEGs by separate.

**Supplemental Table S7.** GOSlim terms and gene lists in vertical format of Spruce over time including gene descriptions analyzing needle-specific, common and root-specific DEGs by separate.

**Supplemental Table S8.** Super cluster Gene Ontology (GO) enrichments and members lists.

**Supplemental Table S9.** Spruce-specific TF-DEC. Details of TF-DEC without any ortholog predicted by Gymno-Plaza.

**Supplemental Table S10.** Functional descriptions of cluster members identified on Transcription Factors differentially regulated by cold (TF-DEC).

**Supplemental Table S11.** Gene Ontology (GO) enrichments on DEGs list over time series of Spruce.

**Supplemental Table S12.** Hubs neighborhood analysis.

**Supplemental Table S13.** Hubs descriptions and topology information.

